# Evidence for gene-environment correlation in child feeding: Links between common genetic variation for BMI in children and parental feeding practices

**DOI:** 10.1101/407221

**Authors:** Saskia Selzam, Tom A. McAdams, Jonathan R. I. Coleman, Susan Carnell, Paul F. O’Reilly, Robert Plomin, Clare H. Llewellyn

## Abstract

The parental feeding practices (PFPs) of excessive restriction of food intake (‘restriction’) and pressure to increase food consumption (‘pressure’) have been argued to causally influence child weight in opposite directions (high restriction causing *overweight*; high pressure causing *underweight*). However child weight could also ‘elicit’ PFPs. A novel approach is to investigate gene-environment correlation between child genetic influences on BMI and PFPs. Genome-wide polygenic scores (GPS) combining BMI-associated variants were created for 10,346 children (including 3,320 DZ twin pairs) from the Twins Early Development Study using results from an independent genome-wide association study meta-analysis. Parental ‘restriction’ and ‘pressure’ were assessed using the Child Feeding Questionnaire. Child BMI standard deviation scores (BMI-SDS) were calculated from children’s height and weight at age 10. Linear regression and fixed family effect models were used to test between-(n=4,445 individuals) and within-family (n=2,164 DZ pairs) associations between the GPS and PFPs. In addition, we performed multivariate twin analyses (n=4,375 twin pairs) to estimate the heritabilities of PFPs and the genetic correlations between BMI-SDS and PFPs. The GPS was correlated with BMI-SDS (β=0.20, *p*=2.41×10^-38^). Consistent with the gene-environment correlation hypothesis, child BMI GPS was positively associated with ‘restriction’ (β=0.05, *p*=4.19×10^-4^), and negatively associated with ‘pressure’ (β=-0.08, *p*=2.70×10^-7^). These results remained consistent after controlling for parental BMI, and after controlling for overall family contributions (within-family analyses). Heritabilities for ‘restriction’ (43% [40-47%]) and ‘pressure’ (54% [50-59%]) were moderate-to-high. Twin-based genetic correlations were moderate and positive between BMI-SDS and ‘restriction’ (r_A_=0.28 [0.23-0.32]), and substantial and negative between BMI-SDS and ‘pressure’ (r_A_=-0.48 [-0.52 --0.44]. Results suggest that the degree to which parents limit or encourage children’s food intake is partly influenced by children’s genetic predispositions to higher or lower BMI. These findings point to an evocative gene-environment correlation in which heritable characteristics in the child elicit parental feeding behaviour.

**Author Summary:** It is widely believed that parents influence their child’s BMI via certain feeding practices. For example, rigid restriction has been argued to cause *overweight*, and pressuring to eat to cause *underweight*. However, recent longitudinal research has not supported this model. An alternative hypothesis is that child BMI, which has a strong genetic basis, evokes parental feeding practices (‘gene-environment correlation’). To test this, we applied two genetic methods in a large sample of 10-year-old children from the Twins Early Development Study: a polygenic score analysis (DNA-based score of common genetic variants robustly associated with BMI in genome-wide meta-analyses), and a twin analysis (comparing resemblance between identical and non-identical twin pairs). Polygenic scores correlated positively with parental restriction of food intake (‘restriction’; β=0.05, *p*=4.19×10^-4^), and negatively with parental pressure to increase food intake (‘pressure’; β=-0.08, *p*=2.70×10^-7^). Associations were unchanged after controlling for all genetic and environmental effects shared within families. Results from twin analyses were consistent. ‘Restriction’ (43%) and ‘pressure’ (54%) were substantially heritable, and a positive genetic correlation between child BMI and ‘restriction’ (*r*_A_=0.28), and negative genetic correlation between child BMI and ‘pressure’ (*r*_A_=-0.48) emerged. These findings challenge the prevailing view that parental behaviours are the sole cause of child BMI by supporting an alternate hypothesis that child BMI also causes parental feeding behaviour.

## Introduction

The home and family environment has been studied for decades with the assumption that it is a crucial determinant of children’s health and development. Since the onset of the childhood obesity crisis at the turn of the century, the spotlight has turned onto environmental factors associated with variation in adiposity, in the hope that modifiable elements may be identified as intervention targets. Perhaps unsurprisingly, parental behaviours have received a great deal of attention. Parents are widely considered to be the ‘gatekeepers’ to their children’s food, and powerful shapers of their developing eating behaviour^1-3^. Two parental feeding practices (PFPs) in particular have been hypothesised to play a causal role in children’s ability to develop good self-regulation of food intake and consequently determine their weight. Excessive restriction of the type and amount of food a child is allowed to eat (‘restriction’) has been hypothesised to lead to overeating when parental restriction is no longer in place, because the child will potentially then hanker after the foods he or she is not usually allowed to eat – the so-called ‘forbidden fruit effect’^1,4,5^. On the other hand, overly pressuring a child to eat, or to finish everything on the plate (‘pressure’), is thought to be anxiety-provoking for a child with a poor appetite, and serves only to increase undereating further, and compromise weight gain^6,7^.

A wealth of cross-sectional findings are consistent with these hypotheses^8^, but another plausible explanation for the observed correlations is that parents are responding to their child’s emerging characteristics, rather than causing them. Parents may only adopt restrictive strategies when a child shows a tendency toward overeating, or gains excessive weight; and they may pressure their child to eat only if he or she is a poor eater, or underweight. The few longitudinal studies testing bidirectionality have shown that children’s weight prospectively predicts PFPs^9-13^. Furthermore, three studies showed no prospective association from PFPs to child weight^10^, and the studies reporting bidirectional relationships found stronger associations from child weight to parental behaviour than the reverse direction^9,11^. Although these findings point towards children’s weight eliciting PFPs, the possibility of residual confounding in observational studies hinders conclusions about causation – temporality does not necessarily mean causality.

Testing whether children genuinely cause their parents’ behaviour presents challenges. It is not possible – practically or ethically – to randomise children to be overweight or underweight, and examine how parents respond. Genetic approaches provide a powerful alternative method of interrogating the role of children in causing their parents’ behaviour towards them, especially for child characteristics with an established genetic basis. To date, no study has applied genetically sensitive methods to test for gene-environment correlation in parental feeding behaviour. Family and twin studies have shown that Body Mass Index (BMI), is highly heritable in both adulthood and late childhood (∼80%)^14-16^. Twin designs can therefore be used to test if parental behaviour has a heritable component, by comparing within-pair resemblance for identical and fraternal twin pairs in childhood. If found, this indicates that parental behaviour is explained to some extent by variation in children’s genotype – termed evocative gene-environment correlation^17^. Twin designs can also be extended to the analysis of multiple variables to establish if genetic influence on a particular child characteristic (e.g. weight) also predicts the parental behaviour of interest (e.g. PFPs). If such analyses show that a child characteristic is genetically correlated with parenting traits, it indicates that these child characteristics influence parenting behaviours. A meta-analysis of 32 twin studies of different types of parenting behaviour reported an average heritability estimate of 23%, indicating that children’s genotype is predictive of a moderate amount of variation in parental behaviour^18^.

Children’s DNA can also be used to test for gene-environment correlation. Genome-wide meta-analyses have made great progress in identifying common single nucleotide polymorphisms (SNPs) that are robustly associated with body mass index (BMI) in adults and children^19^. These can be combined to calculate a genome-wide polygenic score (GPS) that indexes individual-specific propensity to higher or lower BMI, along a continuum, although in the aggregate the GPS explains only a small proportion of variance in BMI (approximately 3%). Nevertheless, children’s BMI GPS can therefore be used to test the hypothesis that parents develop their feeding practices specifically in response to their child’s weight, as indicated by a correlation between child BMI GPS and PFPs. Unlike for other correlations, a possible interpretation for associations between differences in DNA sequence and parental behaviour is genetic causation, because DNA sequence variation cannot be caused by parental behaviour. A caveat to this is that a parent’s feeding practices may reflect their own genetic predisposition to be of a higher or lower BMI, rather than that of their children. In this way, a correlation between child BMI GPS and PFPs may simply reflect a child’s genetic predisposition to be of a higher or lower BMI, which they inherit from their parent with whom they share 50% of their DNA. In addition, genetic effects related to adult BMI discovered in genome-wide association studies could potentially incorporate effects of PFPs if they were to causally influence child BMI, and its trajectory into adulthood. However, within-family designs can circumvent both of these limitations to some extent. Studying variation in PFPs according to variation in BMI GPS within co-twins accounts for both genetic and environmental shared effects within families (e.g. parental genetic predisposition to be of higher or lower BMI). By applying both quantitative and molecular genetic methods, and utilising statistical approaches to account for family effects, we intended to address the various limitations presented by the individual methods.

The goals of this study were to test for gene-environment correlation between children’s BMI and PFPs, using a twin design and a BMI GPS. We hypothesised that: (i) children’s BMI GPS would be positively associated with parental restriction and negatively associated with parental pressure, even after accounting for shared genetic and environmental family influences, and (ii) parental restriction and parental pressure would be moderately heritable, and that genetic influence on PFPs would be partly explained by genetic influence on children’s BMI.

## Results

### Phenotypic correlations

Child BMI-SDS was significantly positively correlated with ‘restriction’ (β = 0.19, *t*(4004) = 12.09, *p* = 4.45×10^-33^, *R*^*2*^ = 0.035), such that parents were more restrictive over their child’s food intake where the child had a higher BMI. In contrast, child BMI-SDS was significantly negatively correlated with ‘pressure’ (β = -0.24, *t*(4058) = -15.59, *p* = 3.14×10^-53^, *R*^*2*^ = 0.056), where parents exerted higher amounts of pressure on their child to eat, if their child was leaner. ‘Restriction’ and ‘pressure’ were significantly positively correlated (β = 0.15, *t*(4207) = 9.51, *p* = 3.08×10^-21^, *R*^*2*^ = 0.021), suggesting that parents who tend to exert higher levels of ‘restriction’ also have a more pressuring feeding style, to some extent.

### Genome-wide Polygenic Score (GPS) analyses

In our sample of unrelated individuals, child BMI GPS was positively correlated with child BMI-SDS (β = 0.20, *t*(4226) = 13.08, *p* = 2.41×10^-38^, *R*^*2*^ = 0.039). Mirroring phenotypic results for child BMI-SDS, children’s BMI GPS was significantly positively correlated with ‘restriction’ (β = 0.05, *t*(4255) = 3.53, *p* = 4.19×10^-4^, *R*^*2*^ = 0.003), and significantly negatively correlated with ‘pressure’ (β = -0.08, *t*(4315) = -5.15, *p* = 2.70×10^-7^, *R*^*2*^ = 0.006) (Fig 1). These findings indicate that children’s genetic predisposition to higher BMI, elicits, to some extent, restrictive feeding behaviours in the parent; whereas children’s genetic predisposition to lower BMI elicits greater pressure to eat by parents.

**Fig 1.**
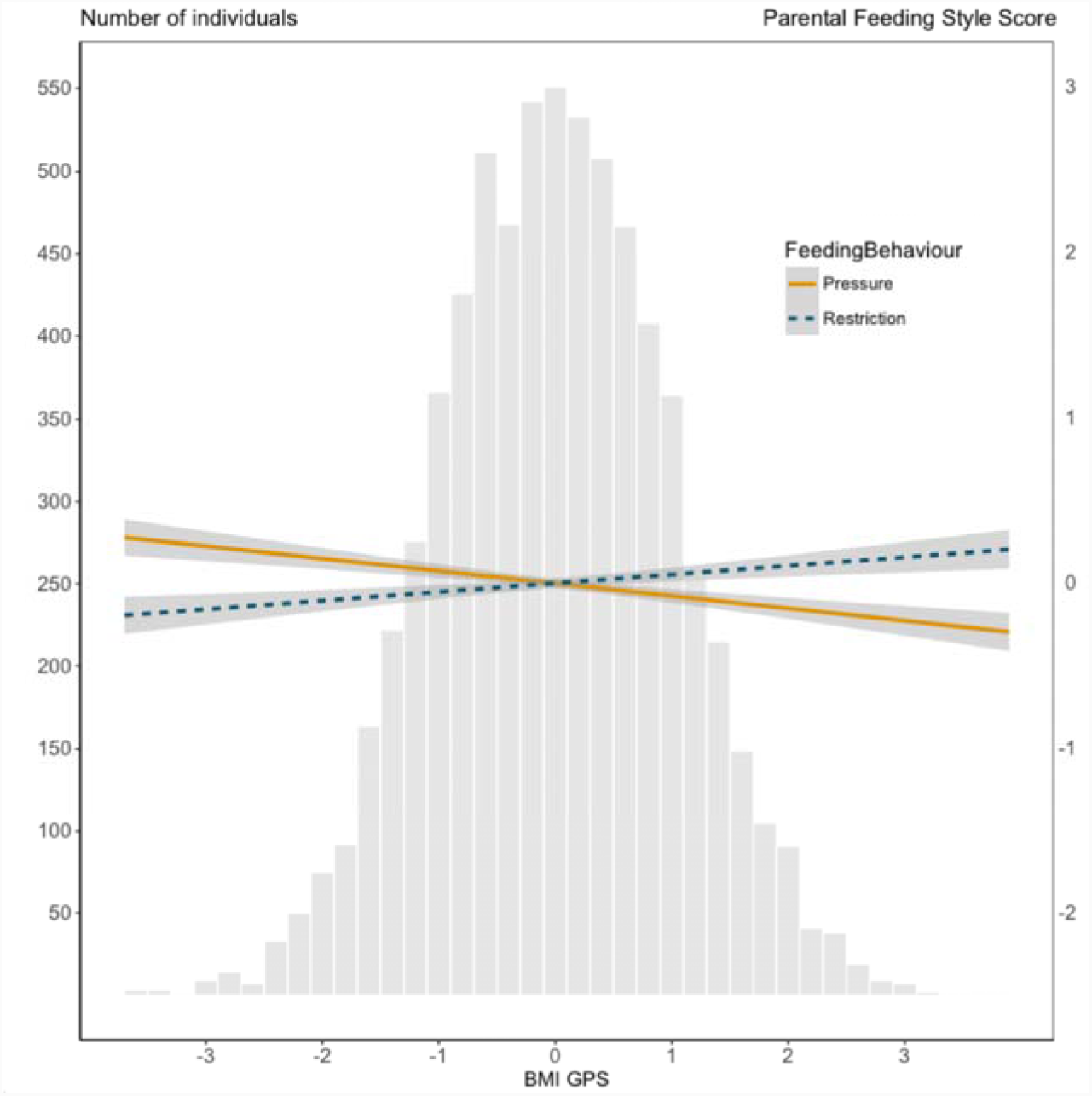
The associations between child BMI polygenic score and parental feeding practices.

Parental BMI correlated positively with child BMI-SDS (β = 0.26, *t*(3761) = 17.00, *p* = 1.57×10^-62^, *R*^*2*^ = 0.071) and ‘restriction’ (β = 0.08, *t*(3711) = 4.64, *p* = 3.65×10^-6^, *R*^2^ = 0.005), but was not significantly associated with ‘pressure’ (β = -0.03, *t*(3757) = -1.68, *p* = 0.09, *R*^*2*^ < 0.001). The magnitude and direction of effects remained identical after controlling for parental BMI in ‘restriction’ (β = 0.05, *t*(3711) = 2.92, *p* = 3.48×10^-3^, *R*^*2*^ = 0.003) and in ‘pressure’ (β = -0.08, *t*(3757) = -4.62, *p* = 3.97×10^-6^, *R*^*2*^ = 0.005).

Child BMI GPS predicting standardized measures of parental ‘restriction’ (β = 0.05, *p* = 4.19×10^-4^) and parental ‘pressure’ (β = -0.08, *p* = 2.70×10^-7^) as indicated by the best-fit regression lines. The grey areas surrounding the best-fit lines represent standard errors of the prediction estimates. The histogram depicts the BMI GPS normal distribution.

### Within-family analyses

To establish the association between children’s BMI GPS and PFPs entirely without confounding by genetic and environmental family factors shared by twin pairs, we performed family fixed effect analyses in DZ co-twins. This analysis examined the extent to which parents vary their ‘restriction’ and ‘pressure’ across twin pairs in response to differences in their BMI GPSs. As shown in Fig 2, beta coefficients for BMI GPS predicting PFPs remained largely stable when comparing unrelated individuals (Model 1) and DZ twin pairs (Model 2). For unrelated individuals (Model child BMI-SDS significantly positively predicted ‘restriction’ and significantly negatively predicted ‘pressure’, as previously reported. The magnitude of the within-family estimates for the combined (same-sex and opposite-sex) DZ co-twins (Model 2) were virtually the same as those for the unrelated individuals for the relationships between BMI GPS and ‘restriction’ (*t*(2054) = 3.50, *p* = 7.10×10^-3^, *Adj. R*^*2*^_*model*_ = 0.724) and BMI GPS and ‘pressure’ (*t*(2103) = -4.82, *p* = 1.52×10^-6^, *Adj. R*^*2*^_*model*_ = 0.641) (*R*^*2*^ magnitudes for Model 2 are large because all shared factors among family members, including genetic and environmental influences, are accounted for). These findings indicate that even when shared family effects are completely accounted for, children’s BMI GPS is significantly associated with PFPs, providing additional evidence that children’s genetic predisposition to BMI evokes certain parental feeding responses. When repeating Model 2 analyses separately for same-sex and opposite-sex DZs, magnitudes of effect sizes (Fig 2) remained consistent for the prediction of ‘pressure’ in same-sex DZ pairs (*t*(1118) = -3.36, *p* = 8.02×10^-4^, *Adj. R*^*2*^_*model*_ = 0.607) and opposite-sex DZ pairs (*t*(984) = -3.49, *p* = 5.12×10^-4^, *Adj. R*^*2*^_*model*_ = 0.678). Although BMI GPS in opposite-sex DZs was a significant predictor of within-family differences in ‘restriction’ (*t*(966) = 3.76, *p* = 1.82×10^-4^, *Adj. R*^*2*^_*model*_ = 0.731), same-sex DZ data did not show a significant within-family association (*t*(1087) = 1.21, *p* = 0.23, *Adj. R*^*2*^_*model*_= 0.719), indicating that within a family environment, GPS differences in BMI between same-sex DZ twins are not related to differences in parental ‘restriction’.

**Fig 2.**
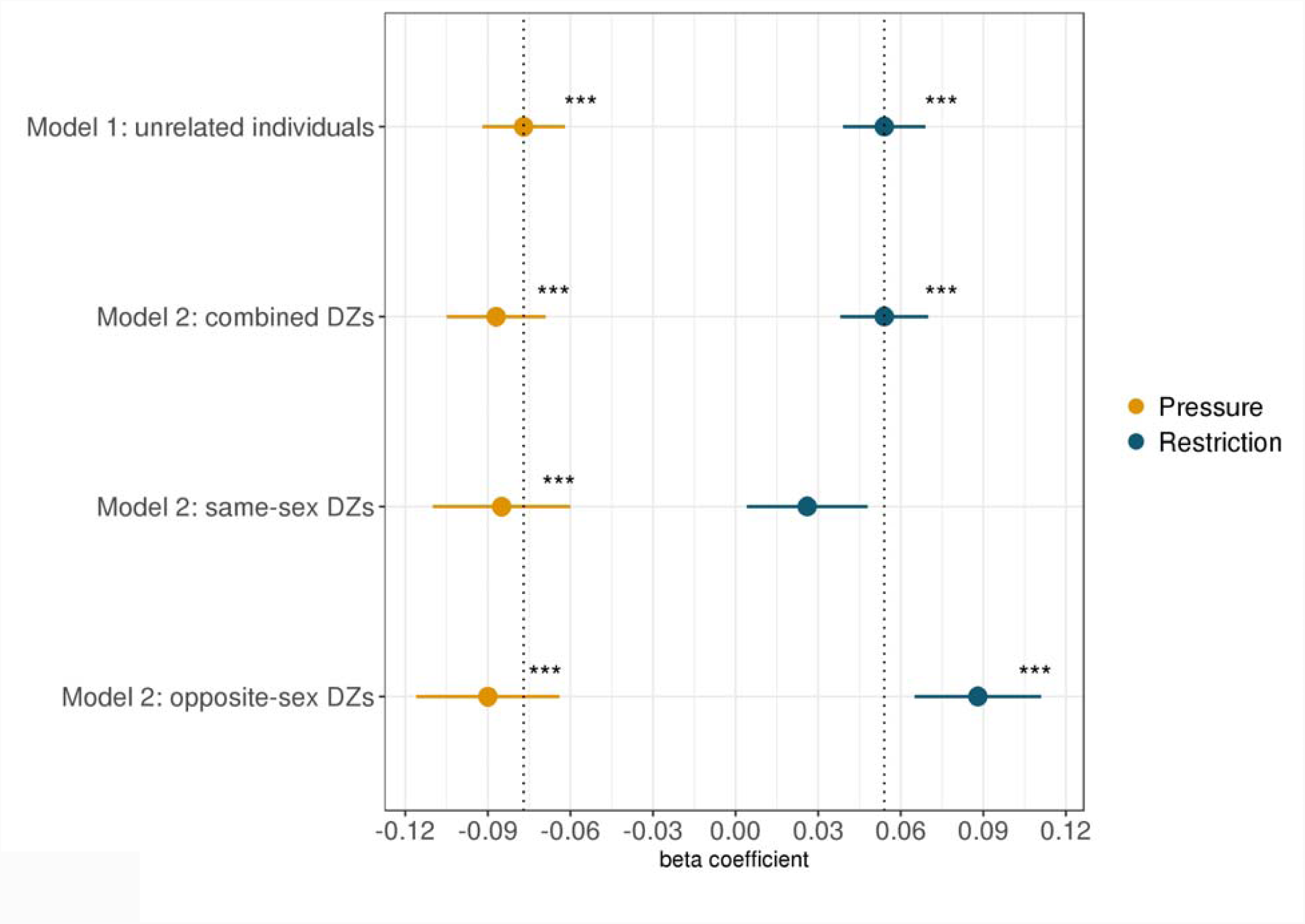
Contrasting results from between-family analyses to results from within-family analyses.

Model 1 describes results using BMI GPS of unrelated individuals to predict PFPs, where β_*GPS*_ indicates the change in the outcome trait per one standard deviation increase in the BMI GPS. Model 2 summarises results using BMI genome-wide polygenic scores in a sample of DZ co-twins using a family fixed effects model, where β_*GPS*_indicates the increase in PFPs within DZ pairs, per one standard deviation increase in BMI GPS within DZ pairs. Model 2 analyses were performed using the combined DZ sample, and same-sex DZ pairs and opposite-sex DZ pairs only. The dotted lines represent the beta coefficient estimates for Model 1. * = *p*<0.05; ** = *p*<0.01; *** = *p*<0.001.

### Twin analyses

We performed multivariate genetic analyses (a correlated factors model) to establish the heritability of ‘restriction’ and ‘pressure’ and to test the extent to which genetic influence on child BMI-SDS elicited PFPs as indicated by the magnitude of genetic correlations between BMI, ‘restriction’, and ‘pressure’. Fig 3 shows the variance components (A, C and E) for each measured phenotype, as well as the genetic, shared environmental and non-shared environmental correlations between phenotypes derived from the correlated factors model (see Supplementary Table S4 for fit statistics and model comparisons, and Supplementary Table S3 for intra-class correlations). Heritability estimates (A) were moderate to high for parental ‘restriction’ (43%, 95% CI [40%, 47%]) and parental ‘pressure’ (54%, 95% CI [50%, 59%]); heritability of child BMI-SDS was high (78%, 95% CI [72%, 84%]). Consistent with the findings from the GPS analyses, there was a significant, positive moderately sized genetic correlation between child BMI-SDS and parental ‘restriction’ (r_A_=0.28, 95% CI [0.23, 0.32]), indicating that some of the genetic effects that predispose a child to a higher BMI also elicit more food restriction by their parent. A sizeable significant negative genetic correlation was observed between child BMI-SDS and parental ‘pressure’ (r_A_=-0.48, 95% CI [-0.52, -0.44]), indicating that many of the genetic effects that predispose a child to a lower BMI elicit greater parental pressure on the child to eat.

**Fig 3.**
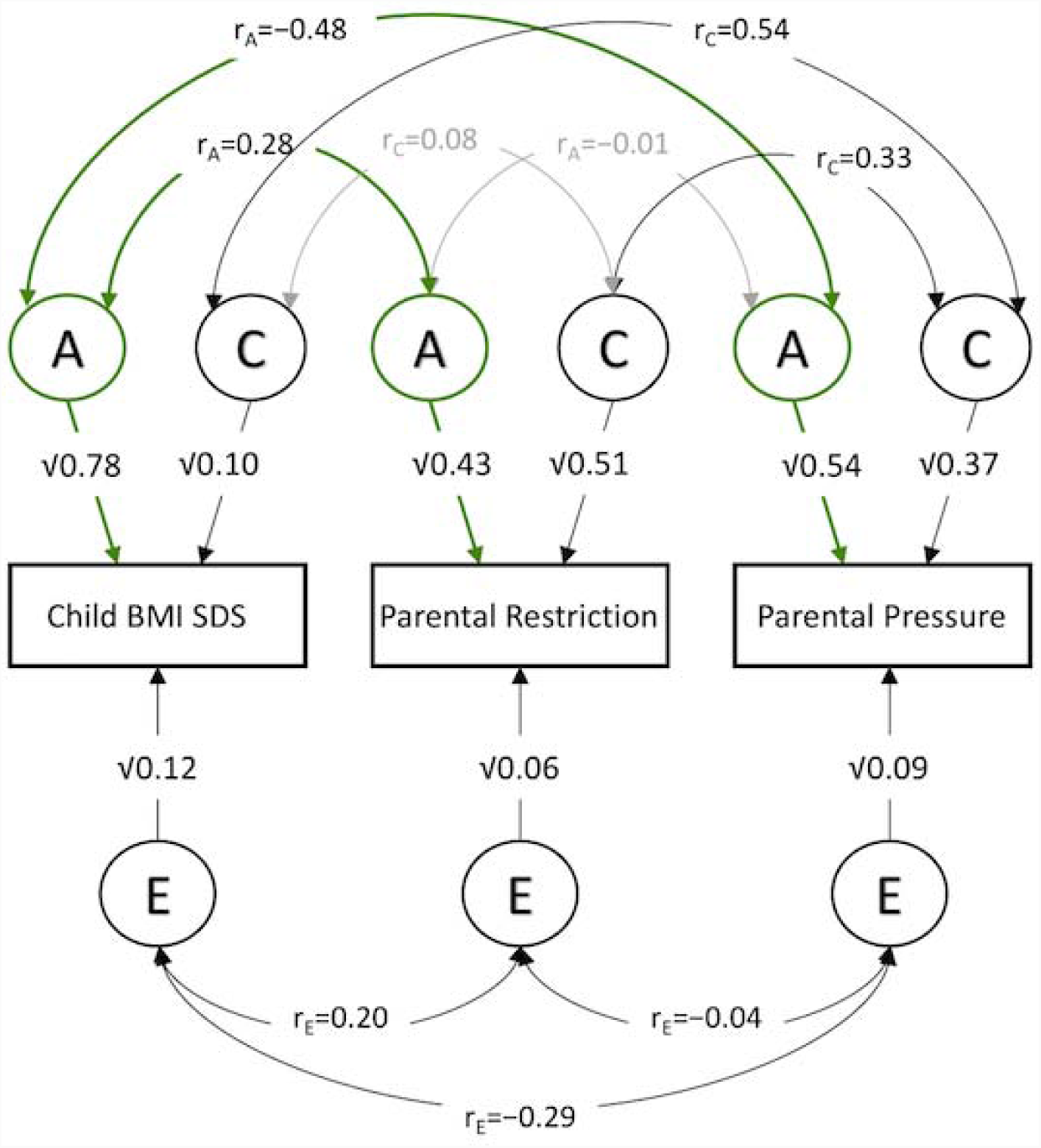
The correlated factors model.

A correlated factors model (males and females combined) showing: (i) the genetic (A), shared environmental (C) and non-shared environmental (E) influences on child BMI SDS, parental restriction and pressure; and (ii) common genetic (r_A_), shared environmental (r_C_) and non-shared environmental (r_E_) correlations between child BMI, and parental restriction and pressure. Grey arrows indicate non-significant associations. Correlations including the 95% confidence intervals can be found in Supplementary Table S5.

## Discussion

### Summary of findings

We describe the first study to test for gene-environment correlation for parental feeding behaviour in relation to child weight, using a twin design and children’s DNA. Results support our hypothesis that parents’ feeding practices are evoked, in part, by their children. Parental ‘restriction’ and ‘pressure’ were positively and negatively associated with child BMI respectively, in keeping with many previous cross-sectional studies^8^. We applied novel genetic methods to show for the first time that children’s BMI GPS was significantly positively associated with ‘restriction’ and negatively associated with ‘pressure’, even after accounting for the potentially confounding shared familial effects (both genetic and environmental). This suggests that children’s genetic influence on weight causally explains part of the observed phenotypic associations. Our twin analysis provided quantitative estimates of the total variance in parental feeding practices explained by children’s genotype. Heritability was substantial for both ‘restriction’ (43%) and ‘pressure’ (54%), indicating that children’s genes explain about half of the variation in parental feeding behaviour. Multivariate twin analysis established the extent to which parental feeding behaviour was determined by children’s genetic influence on BMI *specifically*. The genetic correlations between children’s BMI and both ‘restriction’ (r_A_=0.28) and pressure (r_A_=-0.48) were moderate, indicating overlap between the genes that influence parental feeding behaviour and children’s BMI.

A potential confounder of the association between child GPS and parental feeding behaviour, was the parent’s own genetic propensity to a higher or lower BMI. Children inherit half of each of their parents’ genetic material, so the expected correlation between a child’s GPS with that of their parent’s is 0.50. A parent’s genetic predisposition to be of a higher or lower BMI may also influence the way they feed their children, which could introduce a passive (rather than ‘evocative’) gene-environment correlation. For example, a parent with a higher BMI may be more restrictive over their child’s food intake, but their child also inherits their parent’s susceptibility to be of a higher BMI – restrictive feeding may therefore simply be a marker for a child’s genetic predisposition to be of a higher BMI that is transmitted to them by their parent, rather than a causal risk factor (the same could be true for a more pressuring feeding style and lower BMI). In line with this, parental BMI (indexing parental GPS) was significantly positively associated with parental restriction indicating that parents of a higher weight exert greater restriction over their children’s food intake (β = 0.08); although the association with parental pressure was not significant. Adjustment for parental BMI did not attenuate the associations between child GPS and either restriction or pressure, suggesting it was not confounding the relationship between parental feeding behaviour and child BMI GPS. Nevertheless, adjustment for parental BMI cannot completely remove confounding from parental BMI, nor can it account for the potential effect of longer-term BMI on parental feeding behaviours. However, in order to rule out confounding by any parental characteristics (both genetic and environmental), we took advantage of a family fixed-effect design, which held the effects of family constant while testing the association between the child BMI GPS and parental feeding practices in DZ co-twins. The within-family analysis allowed us to demonstrate that even after accounting for all familial effects, parents vary their feeding behaviour for each child depending on their GPS – larger GPS differences between pairs were associated with more pronounced differences in parental feeding behaviour. The magnitudes of the between-and within-family associations between parental feeding behaviour and child GPS were virtually the same, with the exception of the relationship between child GPS and ‘restriction’ in same-sex twins, strengthening the evidence that children evoke parental responses based on their genetic predispositions for BMI. Nevertheless, as expected and in keeping with the small amount of variance in explained in BMI by the GPS, the size of the associations between the BMI GPS and PFPs were small.

### Other relevant research

The findings from this study accord with those from twin studies of many other types of parenting behaviours that have also tended to show moderate heritability. A meta-analysis of 32 child twin studies on maternal positivity, negativity, affect and control in relation to parenting showed an average heritability of 24%^18^, indicating widespread, child-driven genetic influences on parental behaviour. The heritability estimates for ‘restriction’ (43%) and ‘pressure’ (54%) were somewhat higher than the average heritability estimate for the parenting styles considered in the meta-analysis (24%), but in keeping with the magnitude of the heritability of negative parenting styles observed across early childhood (∼55%)^20^.

Despite providing evidence for gene-environment correlation, the design of our study was not able to shed light on the reverse causal direction – the influence of PFPs on child weight. The few prospective studies that have attempted to establish the cause-effect relationship in the parent-child dynamic using bidirectional analyses have suggested either only a small effect of restriction and/or pressure on child weight, or none^9-11,13^. Prospective studies therefore suggest that PFPs may be less important than is commonly assumed. The well-established strong genetic influence on children’s weight – in the order of 70-80%^15,16^ – also supports the hypothesis that parents influence child weight via genetic inheritance more than by creating an ‘obesogenic’ family environment. However, it cannot be ruled out that genetic effects related to BMI in the parents also contribute to an obesogenic environment if gene-environment correlation was at play, further passively reinforcing the child’s inherited genetic propensities. The shared environmental influence on BMI in late childhood is also low^15,16^. In the current study, the shared environmental influence on parental feeding behaviour was the proportion of variance that was common to both twins in a pair (invariant within families). It therefore likely reflects variation in feeding behaviour that was parent-driven rather than child-directed. These estimates indicated that a substantial proportion of variation in both ‘restriction’ (C=43%) and ‘pressure’ (C=37%) were also parent-origin.

Experimental studies in the form of large well-designed randomised controlled trials (RCTs) are needed to truly test the hypothesis that PFPs causally modify children’s weight gain trajectories. Very few of these have been conducted to date, and they have focused on the preschool years. Nevertheless, two landmark studies have indicated that parental behaviour may, in fact, be influential in early life. NOURISH^21^ was an Australian RCT that randomised 352 parents and infants to receive a feeding intervention (including using low amounts of pressure, and employing child-responsive methods of food restriction) during the period of complementary feeding; 346 families were randomised to the standard care control group. At three to four years of age, children in the intervention group had better appetite control than those in the control group, and there were fewer overweight children; although this did not reach statistical significance^22^. INSIGHT^23^, a US RCT, randomised 145 new mothers to a responsive parenting intervention that focused on feeding infants only in response to their hunger and satiety signals (but neither pressuring nor restricting their milk and food intake), during milk-feeding and complementary feeding; 145 mothers were randomised to a control group. At one year significantly fewer infants in the intervention group were overweight (6%) compared to the control group (13%). These RCTs indicate that parental feeding behaviour can modify young children’s eating behaviour and weight gain. However, these studies were conducted in infants and young preschool children so it is unclear whether these findings are generalisable to older children.

The genetic correlations between children’s BMI and parental feeding behaviour were modest, and were far from complete (i.e. less than 1.0), indicating that other genetically-determined child characteristics are also influencing parental feeding behaviour. Children’s appetite is under strong genetic control; twin studies – including this sample – have shown high heritability for appetite^24,25^ and shared heritability with BMI^26^, and appetite is associated with the BMI GPS in this sample and has been shown to mediate part of the GPS-BMI association^27^. It is therefore likely that child appetite also influences parental feeding behaviour^24,25^. In support of this, prospective and within-family studies have provided evidence that within the context of parental feeding, parents respond not only to their child’s weight but to their eating styles too. A large prospective population-based study used bidirectional analyses to show that parents whose children were excessively fussy at baseline increased their pressure over time^28^. A reverse relationship also pertained, but the temporal association from child to parent was stronger. A large within-family study of preschool twins showed that parents varied their pressuring feeding style when their twins were discordant for food fussiness^29^. The fussier twin was pressured more than their co-twin, also in support of a child-driven model of parental feeding behaviour. It stands to reason that a child who is a picky eater is pressured, to try some of their vegetables or to eat more overall. Along the same lines, a natural response from a parent who has a child who shows a tendency toward excess intake and a relatively pronounced preference for foods rich in sugar or fat, is to enforce some restriction.

We also found a positive phenotypic correlation between ‘restriction’ and ‘pressure’ (β = 0.15), indicating that parents who exert higher levels of restriction on their children also tend to pressure them more. This suggests that some parents have a more controlling feeding style in general.

### Implications and future research

The relationship between parental behaviour and children’s emerging characteristics appears to be reciprocal and complex. The current findings suggests that parents’ feeding responses to child weight are to exert greater restriction of food intake on children with a higher BMI, and to pressure a thinner child to eat. However, these strategies may not be effective in the long run. RCTs have suggested that PFPs can have a lasting and important impact on children’s weight and eating behaviour in the early years, although whether or not these findings apply to older children has yet to be determined. It is well established that the genetic influence on the BMI in younger children is lower, and the shared environmental effect is higher, than in older children^15,16^. This suggests that parental influence diminishes as children grow older, gain independence and spend increasing time outside the home with peers rather than parents^30^. Large RCTs that follow children from early life to later childhood are needed to establish if PFPs influence the weight of older children.

### Strengths & Limitations

A strength of this study is that we used several genetically sensitive methodological approaches to explore the causal relationships between child BMI and PFPs, yielding consistent results. PFPs were measured using the Child Feeding Questionnaire, which has well established criterion and construct validity, as well as good internal and test-retest reliability^31^. This instrument has been used widely in previous research into child weight, allowing the findings from this study to be directly compared to a wealth of existing results.

A potential limitation is that heritability estimates from twin studies rely on the assumption that MZs and DZs share their environment in terms of the trait in question to the same extent, so-called the ‘equal environments assumption’; if this is violated, the findings are invalid. Therefore if parents feed MZs more similarly than DZs simply because they are identical, this would artificially inflate the MZ correlation and, consequently, heritability. However, if MZs are fed more similarly than DZs because parents are responding to their genetically determined BMI or traits that share genetic influence with BMI such as appetite, differences in feeding experience across MZs and DZs do not constitute a violation of the equal environments assumption because these differences in feeding practices are being driven by greater genetic similarity between MZs than DZs. In addition, if parents’ reports of how similarly they fed their twins were biased by their perceived zygosity (i.e. reported treatment was not a true reflection of actual treatment, but related to the twins being MZ or DZ), this would also render the heritability estimates unreliable.

However, this seems unlikely given previous findings that parents’ reports about their twins’ are not biased by their beliefs about their zygosity, using the ‘mistaken zygosity’ design^32^.

Another limitation was the lack of parental genotypes assessments. Parental BMI is by no means a perfect proxy for their genotypic predisposition to higher or lower BMI; the most powerful approach would be to have parental genotypes whereby the non-transmitted alleles from the parents (which relate to their own BMI and behaviour, but not to that of their child) can be entirely separated from the child’s genotype. Nevertheless, the within-family analysis controlled for all family-level genetic and environmental effects, and the magnitudes of the relationships between child BMI and PFPs were unaffected. A further limitation is that we were unable to validate self-reported parental BMI, which may have been inaccurate and could potentially bias our results. Additionally, it may be possible that PFPs are largely explained by environmental factors that influence children’s BMI. As the BMI GPS is not yet strong enough to be a sufficient proxy to separate genetic and environmental effects on child BMI, we were unable to test this question empirically. However, considerable genetic correlations between child BMI and PFPs derived from the twin model renders this explanation unlikely. Lastly, BMI was only reported at one time point, but PFPs are likely to be driven by the child’s emerging BMI throughout the developmental years. However, BMI-associated SNPs and BMI GPS are associated with weight gain trajectories from infancy throughout childhood, so the BMI GPS in fact captures a long window of child BMI^14,33^.

### Conclusion

This study provides new evidence for gene-environment correlation in parental feeding practices. We have shown that parental feeding practices are substantially heritable and are elicited by the genes that influence children’s BMI. Genome-wide polygenic scores that index children’s genetic propensities for their BMI significantly predicted their parents’ feeding practices, even after potentially confounding shared family effects were taken into account. The findings of this study provide a new perspective on the nature of the associations between parental feeding practices and child BMI.

## Methods

### Sample

Participants were from the Twins Early Development Study (TEDS). Between 1994-1996 TEDS recruited over 15,000 twin pairs born in England and Wales, who have been assessed in multiple waves across their development up until the present date. Despite some attrition, about 10,000 twin pairs still actively contribute to TEDS, providing genetic, cognitive, psychological and behavioural data. TEDS participants and their families are representative of families in the UK^34^. Written informed consent was obtained from parents before data collection began. Project approval was granted by King’s College London’s ethics committee for the Institute of Psychiatry, Psychology and Neuroscience (05.Q0706/228). This study included 4,445 unrelated individuals with genotyping for the GPS analysis, 2,164 genotyped dizygotic (DZ) twin pairs (1,151 same-sex DZ pairs, 1,013 opposite-sex DZ pairs), and 4,375 twin pairs for the twin analysis (1,636 monozygotic (MZ) pairs, 1,441 same-sex DZ pairs, and 1,298 opposite-sex DZ pairs).

### Genotyping

Two different genotyping platforms were used because genotyping was undertaken in two separate waves, five years apart. AffymetrixGeneChip 6.0 SNP arrays were used to genotype 3,665 individuals at Affymetrix, Santa Clara (California, USA) based on buccal cell DNA samples. Genotypes were generated at the Wellcome Trust Sanger Institute (Hinxton, UK) as part of the Wellcome Trust Case Control Consortium 2 (https://www.wtccc.org.uk/ccc2/). Additionally, 8,122 individuals (including 3,607 dizygotic co-twin samples) were genotyped on HumanOmniExpressExome-8v1.2 arrays at the Molecular Genetics Laboratories of the Medical Research Council Social, Genetic Developmental Psychiatry Centre, using DNA that was extracted from saliva samples. After quality control, 635,269 SNPs remained for AffymetrixGeneChip 6.0 genotypes, and 559,772 SNPs for HumanOmniExpressExome genotypes.

Genotypes from the two platforms were separately phased using EAGLE2^35^, and imputed into the Haplotype Reference Consortium (release 1.1) through the Sanger Imputation Service03/09/2018 06:49:00 before merging genotype data from both platforms. Genotypes from a total of 10,346 samples (including 3,320 dizygotic twin pairs and 7,026 unrelated individuals) passed quality control, including 3,057 individuals genotyped on Affymetrix and 7,289 individuals genotyped on Illumina. The final data contained 7,363,646 genotyped or well imputed SNPs (for more details, see Supplementary Methods S1).

We performed principal component analysis on a subset of 39,353 common (MAF > 5%), perfectly imputed (info = 1) autosomal SNPs, after stringent pruning to remove markers in linkage disequilibrium (r^2^ > 0.1) and excluding high linkage disequilibrium genomic regions so as to ensure that only genome-wide effects were detected.

### Phenotypic measures

The samples used for the analyses differed by necessity in order to accommodate the different methodological approaches: unrelated genotyped individuals (UG); dizygotic genotyped co-twins (DG); twin sample (TS) for quantitative genetic analysis. For the classical twin model approach, only phenotypic data from genotyped twins and their co-twins were selected for comparability across the study samples. Descriptive statistics for all phenotypic measures are reported in Supplementary Table S1a for unrelated genotyped individuals, in Supplementary Table S1b for genotyped DZ twin pairs and in Supplementary Table S1c for samples used for twin modelling.

Children’s body mass index (BMI) was calculated from parent-reported weight (kg) divided by the square of parent-reported height (metres): kg/m^2^. The 1990 UK growth reference data^36^ were used to create BMI standard deviation scores (BMI-SDS) which take account of the child’s age and sex, and represent the difference between a child’s BMI and the mean BMI of the reference children of the same age and sex. BMI-SDS are used rather than BMI itself because BMI varies substantially by age and sex until early adulthood. Reference BMI-SDS have a mean of 0 and a SD of 1: a value greater than 0 indicates a higher BMI than the mean in 1990; a value less than 0 indicates a lower BMI than the mean in 1990. The validity of parent-reported height and weight was tested through home-visits of researchers in a subset of 228 families. Correlations between measurements taken by parents and researchers were high (height: *r* = 0.90; weight: *r* = 0.83) ^37^. BMI-SDS were available for 4,259 (UG), 4,134 (DG), and 8,406 (TS) individuals. Children had a mean age of 9.91 years (*SD*=0.87) when anthropometric measures were assessed.

Parental BMI was calculated for 4,112 individuals using self-reported weight (kg) and height (metres) of the responding parent (kg/m^2^), which was assessed at the same time as childhood height and weight. To account for the gender of the responding parent (97% mothers, 3% fathers), we used the z-standardized residuals of gender-corrected BMI in analyses.

To assess PFPs, we used the Child Feeding Questionnaire^38^, which parents completed when their twins were approximately 10 years old (mean=9.91 years, *SD*=0.87). To measure the degree to which parents restricted their children’s food intake (‘restriction’), we calculated a mean composite score based on 6 items (Cronbach’s alpha = 0.78), such as “I intentionally keep some foods out of my child’s reach“, or “If I did not guide my child’s eating, he/she would eat too many junk foods”. Data were available for 4,386 (UG), 4,228 (DG) and 8,582 (TS) children. Similarly, we created a mean composite score to assess the amount of pressure parents exerted on their children to increase their food intake (‘pressure’), including 4 items (Cronbach’s alpha = 0.61) such as “If my child says “I’m not hungry”, I try to get him/her to eat anyway”, or “I have to be especially careful to make sure my child eats enough”. Data were available for 4,445 (UG), 4,328 (DG) and 8,750 (TS) children. All items were scored on a 5-point Likert scale (Disagree, Slightly disagree, Neutral, Slightly agree, Agree).

### Phenotypic exclusions

For child and parent anthropometrics we removed extreme outliers with implausible values that were deemed to be errors. For children we excluded values based on the following criteria: -/+ 5 standard deviations above or below the mean of height SDS, weight SDS or BMI-SDS; shorter than 105 cm or taller than 180cm; lighter than 12 kg or heavier than 80 kg. After removing outliers, child BMI-SDS had a mean of 0 and a standard deviation of 0.99, showing that the sample is representative of the UK reference population for BMI in 1990 (mean = 0; SD = 1). For parental BMI, we removed individuals with values that fell -/+ 3.5 standard deviations above or below the mean, as well as individuals that weighed below 35 kg. To account for the positive skew, we log transformed this variable. As all variables showed age or sex effects (described in Supplementary Table S1a, S1b, S1c), we controlled for these variables by applying the regression method, using z-standardized residuals for all further analyses. Supplementary Table S2a, S2b and S2c show descriptive statistics for all clean measures (regressed onto age and sex) in unrelated samples, for DZ twin pair samples, and individuals used for twin modelling, respectively.

### Genotypic measures

We created Genome-wide Polygenic Scores (GPSs) for BMI, using summary statistics of the most powerful published genome-wide meta-analysis of BMI to date of 339,224 participants^19^. We calculated a GPS for each individual as the sum of the weighted count of BMI-increasing alleles:

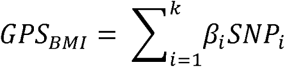

where *i* ∈ {1,2,‥,*k*} and indexes *SNP*_*i*_ and the *i* number of the k BMI increasing alleles included in the score is determined by the *p*-value threshold of the SNP– phenotype association in the discovery GWAS, the *β*-coefficients for each respective genetic variant is used as a weight, and the count of each reference allele is represented by genotype dosage (0,1, or 2 alleles) of *SNP*_*i*_.

We used the software PRSice^39^ to calculate GPS in our sample. To account for multicollinearity among SNPs in Linkage Disequilibrium (LD), which can upwardly bias GPS predictions^40^, genome-wide clumping was performed (r^2^ = 0.1, kb = 250). Using the clumped, independent SNPs, we created eight GPS for 10,346 individuals (7,026 unrelated individuals; 3,320 DZ twin pairs) using increasingly liberal GWAS p-value thresholds (pT: 0.001,0.05,0.1,0.2,0.3,0.4,0.5,1). Diagonals in Supplementary Fig S1 show the number of SNPs included in each respective GPS. As all thresholds performed similarly well (Supplementary Fig S1), we used a GPS based on the smallest *p*-value threshold of 0.001 for all further analyses. Potential effects due to population stratification and genotyping were accounted for by regressing the first ten principal components, and factors capturing genotyping information (microarray, batch and plate) onto the child BMI GPSs, subsequently using the z-standardised residuals in our analyses.

## Statistical Analysis

### Genome-wide Polygenic Score (GPS) analyses

*Trait prediction in unrelated samples*

Associations between child BMI GPS and phenotypes were assessed using linear regression analyses. All variables were standardised prior to analyses, therefore β coefficients from linear regression models are equivalent to Pearson’s correlation coefficients.

*Accounting for family effects in unrelated samples and DZ twin pairs*

Children not only inherit half of each of their parent’s DNA, but also the family environment. Therefore, it is possible that familial effects, both genetic and environmental, confound the relationships between child GPS and PFPs. To account for these potential confounding effects, we used two approaches. Firstly, we removed variance in the PFPs (restriction, pressure) explained by parental BMI in our sample of unrelated individuals using the regression method, and repeated association analyses. Secondly, we used data on genotyped DZ twin pairs to explicitly model the effect of within-DZ twin pair GPS differences on differences in PFPs by accounting for the family contributions in a fixed-effects model:

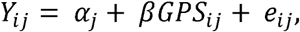

where i E {1,2} indexes the individuals of the dizygotic twin pairs, and j ∈ {1,2,‥,k} indexes the k families (i.e. sets of dizygotic twin pairs). Thus, Y_ij_ is the trait value for the ith individual of the jth family, α_j_ is a vector including the (fixed) family effects, β is the effect of the GPS within families, e_ij_ is the random error for each individual and each family with e_ij_ ∼ N(0, σ^2^), and Cov(α_j_,e _ij_) = 0. The family units were coded using dummy variables in order to estimate the α_j_ effects. By accounting for the differences in contributing factors between families via α_j_, this model tests for the effect of differences in GPS values between DZ twins on the outcome and therefore assesses the impact of GPS with shared genetic and shared environmental factors accounted for. The within-family associations indicate the extent to which parents vary their ‘restriction’ or ‘pressure’ in response to differences in their twins’ BMI GPS. A larger association indicates that the greater the difference between twin pairs’ BMI GPS, the greater the difference in parental ‘restriction’ or ‘pressure’ across two twins in a pair. We applied fixed-effects models to our combined DZ data, and repeated these analyses using same-sex DZ pairs and opposite-sex DZ samples only.

### Twin modelling

To obtain broad estimates of the extent to which individual differences in PFPs are determined by children’s genotypes, we used a multivariate ‘correlated factors’ twin model. This allowed us to estimate: (1) the heritability of PFPs, which provided an indication of the extent to which PFPs are caused by children’s genotypes in general; and (2) the extent of *common* genetic influence on both child BMI-SDS and PFPs, which provided an indication of the extent to which PFPs are caused by children’s genetic propensity to higher or lower BMI, specifically.

Based on biometrical genetics theory^41^, it is possible to decompose variance in a single trait into three components: additive genetic (A; heritability), shared environmental (C; all environmental effects that make family members more similar) and non-shared environmental (E; all environmental effects that contribute to dissimilarities across family members, including random error measurement). The basis of the method is to compare resemblance for a single trait between monozygotic (MZ) and dizygotic (DZ) twin pairs, who share 100% and 50% (on average) of their segregating genetic material, respectively, while both types of twins are correlated 100% for their shared environmental influence. The observed covariation between MZ and DZ pairs is compared with the expected covariation, based on the knowledge of different degrees of allele sharing (or identity by descent (IBD)) of MZ (IBD = 1.0) and DZ pairs (IBD = 0.5 on average). The twin method therefore assumes that MZ and DZ twins share their environments in terms of the trait in question to the same extent (so-called the ‘equal environments assumption’), and the only difference between the two types of twins is the extent of their genetic relatedness.

Using the same principles, comparison of MZ and DZ covariation *across traits* (so-called cross-twin cross-trait covariance, e.g. the covariation between Twin 1 BMI-SDS and Twin 2 ‘restriction’) provides an indication of the extent to which the genetic and environmental influences on multiple traits are the same. The key pieces of information provided are the aetiological correlations, which indicate the extent to which child BMI and PFPs are caused by the same additive genetic (‘genetic correlation’, r_A_), shared environmental (‘shared environmental correlation’, r_C_), and non-shared environmental influences (‘non-shared environmental correlation’, r_E_). In this analysis we were primarily interested in the genetic correlation, which indicates the extent to which the additive genetic influences on child BMI cause PFPs. The aetiological correlations range from -1 to 1 and can be interpreted similarly to Pearson’s correlations. For example, a high *positive* genetic correlation between ‘restriction’ and BMI would indicate that many of the DNA variants that cause higher child BMI are the same as those cause *higher* levels of ‘restriction’, while a high *negative* genetic correlation would indicate that many of the DNA variants causing higher child BMI are the same as those causing *lower* levels of ‘restriction’.

Maximum likelihood structural equation modelling was used to estimate intra-class correlations across the zygosities, the A, C and E parameter estimates and aetiological correlations (with 95% confidence intervals), and goodness-of-fit statistics. Sex differences in the parameter estimates were also tested for using a sex-limitation model. Analyses were implemented in the R package *OpenMx*^42^.

## Author Contributions

Study concept and design: SS, RP, CHL. Processed and quality controlled genotype data: SS. Supervision of genotype quality control: JC. Analysis of data: SS. Interpretation of data: SS, TAM, RP, CHL. Wrote the paper: SS, CHL. Contributed to and critically reviewed the manuscript: All authors

